# Cannabis-associated symptom profiles in patients with first episode psychosis and population controls

**DOI:** 10.1101/577932

**Authors:** Diego Quattrone, Laura Ferraro, Giada Tripoli, Erika La Cascia, Harriet Quigley, EU-GEI group, Charlotte Gayer-Anderson, Peter B. Jones, James B. Kirkbride, Daniele La Barbera, Ilaria Tarricone, Michael T. Lynskey, Tom P. Freeman, Pak C. Sham, Alaistar Cardno, Evangelos Vassos, Jim van Os, Craig Morgan, Ulrich Reininghaus, Cathryn M. Lewis, Robin M. Murray, Marta Di Forti

## Abstract

**Objective:** The evidence is mixed on whether cannabis use is associated with a particular symptomatology in first episode psychosis (FEP) patients.

The authors set out to investigate a) patterns of association between cannabis use and transdiagnostic symptom dimensions; b) whether the extent of use of cannabis contributes to the variation in clinical and subclinical symptom profiles.

**Method:** The authors analysed data from 901 patients and 1235 controls recruited across six countries, as part of the European Network of National Schizophrenia Networks Studying Gene-Environment Interactions (EU-GEI) study. Item response modelling was used to estimate two bifactor models, which included general and specific dimensions of psychotic symptoms in patients and psychotic experiences in controls. The associations between these dimensions and cannabis use was evaluated using linear mixed effects models analyses.

**Results:** In patients, there was a linear relationship between the positive symptom dimension and the extent of lifetime exposure to cannabis, with daily users of high potency cannabis having the highest score (B=0.35; 95%CI 0.14 to 0.56). Moreover, negative symptoms were more common among patients who never used cannabis compared with those with any pattern of use (B=-0.27; 95%CI −0.42 to −0.12).

In controls, psychotic experiences were associated with current use of cannabis but not with the extent of lifetime use.

Neither patients nor controls presented differences in the depressive dimension related to cannabis use.

**Conclusions:** The extent of use of cannabis explains part of the heterogeneous distribution of positive and negative symptoms of FEP patients.

## Introduction

There is compelling evidence to suggest an association between cannabis use and psychotic disorders, with highest risk among those using high potency cannabis on a daily basis (1). However, how cannabis use affects the symptoms psychotic patients present with remains unclear.

The existence of psychotic symptomatology particularly associated with cannabis use has been historically described in several case series (2–5). Nevertheless, case and cohort studies have found mixed results as to whether (6–11) or not (12–15) psychotic patients using cannabis present with more positive symptoms than those not using cannabis. Moreover, there is limited and mixed evidence of any relationship between cannabis use and negative symptoms in psychosis. Some reports suggest fewer negative symptoms in psychotic patients that use of cannabis (16, 17), which is consistent with having enough social skills to obtain the substance. However, this association has not been confirmed in other studies (7, 8) and others even reported a positive association (9).

These inconsistencies might be explained by differences in study design and methods. For example, only a few findings were based on first episode psychosis (FEP) patients (7, 8, 10), which allows one to reduce selection and recall bias, set a common time point and minimise the confounding effect of antipsychotic drugs on symptoms. In addition, a metanalysis of longitudinal studies concluded that most results lacked sufficient power to detect an effect of cannabis on symptoms, or inadequately controlled for potential confounders (18). Furthermore, although a few studies have information on frequency of use, all failed to obtain detailed information on the lifetime pattern of cannabis use, especially on the type and strength of cannabis used. Of note, potent cannabis varieties, with high concentrations of Delta-9-Tetrahyđrocannabinol (Δ9-THC), have been associated with the most harm to mental health (19, 20) and, in recent years, they have become more available worldwide (21, 22). Finally, although the use of a symptom-based approach would be an essential requirement for understanding to what extent environmental factors influence the clinical heterogeneity of psychosis (23), no studies have investigated how cannabis use is related to different psychotic symptom dimensions.

On the other hand, in the general population there are consistent findings regarding the association between cannabis use and subthreshold psychotic symptoms (24). However, most studies presented bias in the selection of non-psychotic population, and all had limited geographical coverage (24).

In this study, we set out to clarify the association between detailed patterns of cannabis use and transdiagnostic symptom dimensions in a large multinational FEP sample.

In addition, we examine the association between detailed patterns of cannabis use and subclinical symptom dimensions in a large sample of controls, that are representative of the population at risk in each catchment area.

Specifically, we sought to test the hypotheses that: (1) positive psychotic symptoms are more common among FEP patients with more frequent lifetime use of cannabis and greater exposure to use of high potency varieties; (2) positive psychotic experiences are more common in population controls with a recent use of cannabis; (3) negative symptoms are more common among those patients who have never used cannabis.

## Methods

### Study design and participants

This analysis is based on data from the incidence and case-control study work package of the European network of national schizophrenia networks studying Gene-Environment Interactions (EU-GEI).

FEP individuals were identified between 2010 and 2015 across six countries to examine incidence rates of schizophrenia and other psychotic disorders (25), and symptomatology at psychosis onset (23). For examining risk factors, we sought to perform an extensive assessment on approximately 1,000 FEP patients and 1,000 population-based controls during the same time period.

Patients were included in the case-control study if they met the following criteria during the recruitment period: (a) aged between 18 and 64 years; (b) presentation with a clinical diagnosis for an untreated FEP, even if longstanding [International Statistical Classification of Diseases and Related Health Problems, Tenth Revision (ICD-10) codes F20-F33]; (c) resident within the catchment area. Exclusion criteria were: (a) previous contact with psychiatric services for psychosis; (b) psychotic symptoms with any evidence of organic causation; and (c) transient psychotic symptoms resulting from acute intoxication (ICD-10: F1x.5).

The recruitment of controls followed a mixture of random and quota sampling methods, in order to achieve the best possible representativeness in age, sex, and ethnicity of the population living in each catchment area. The identification process varied by site and was based on locally available sampling frames, including mostly the use of lists of all postal addresses and general practitioners’ lists from randomly selected surgeries. When these resources were not fully available, internet and newspapers advertising were used to fill quotas. Exclusion criteria for controls were: (a) diagnosis of a psychotic disorder; (b) ever having been treated for psychotic symptoms.

We analysed data from eleven catchment areas, including urban and less urban populations (i.e. Southeast London, Cambridgeshire and Peterborough (England); central Amsterdam, Gouda and Voorhout (the Netherlands); Bologna municipality, city of Palermo (Italy); Paris [Val-de-Marne], Puy-de-Dôme (France); Madrid [Vallecas], Barcelona (Spain); and Ribeirão Preto (Brazil) (23). Further information on the case-control sample and the recruitment strategies is included in the supplementary material.

### Measures

Data on age, sex, and ethnicity was collected using a modified version of the Medical Research Council Sociodemographic Schedule (26). The Operational CRITeria (OPCRIT) system (27) was used by centrally trained investigators, whose reliability was assessed before and throughout the study (k=0.7), to assess psychopathology and generate research-based diagnoses based on different diagnostic classification systems. The Community Assessment of Psychic Experiences (CAPE) (28) was administered to controls to self-report their psychotic experiences. The reliability of the CAPE is good for all the languages spoken in the countries forming part of the EU-GEI study (http://cape42.homestead.com).

A modified version of the Cannabis Experience Questionnaire (CEQ_EU-GEI_) (29) was used to collect extensive information on the patterns of use of cannabis and other drugs. Based on our EU-GEI paper on cannabis (30) we included six measures of cannabis use (Supplementary Table S2), along with a variable measuring specific patterns of cannabis exposure by combining the frequency of use with the potency of cannabis. As illustrated in the supplementary material, the cannabis potency variable was based on the data published in the EMCDDA 2016 (31, 32).

We selected confounders based on their possible association with cannabis use and/or symptom dimensions. These included: sex; age; ethnicity; use of stimulants, hallucinogens, ketamine, cocaine, crack, and novel psychoactive substances (current use no=o; ves=1); current use of cigarettes (no use or smoking less than 10 cigarettes per đav=o; smoking 10 cigarettes or more per day=1), and current use of alcohol (drinking less than 10 units per week= 0; drinking more than 10 alcohol units per week=1).

## Statistical analysis

### Dimensions of psychotic symptoms in patients and psychotic experiences in controls

Data from OPCRIT and CAPE were analysed using multidimensional item response modelling in M*plus*, version 7.4 (33), to estimate two bifactor models, based on the associations among observer ratings of psychotic symptoms in patients and self ratings of psychotic experiences in controls. This methodology is described in full in our EU-GEI paper on symptom dimensions in FEP patients (30), and it was likewise applied to psychotic experiences in population controls. Briefly, CAPE items were dichotomized as 0 ‘absent’ or 1 ‘present’. In order to ensure sufficient covariance coverage for item response modelling, we used items with a valid frequency of ‘present’ ≥10% in our sample, and we excluded those items with low correlation values (<.3) based on the examination of the item correlation matrix. Data used in the analysis contained missing values, which we assumed to be missing at random, allowing for the maximum likelihood estimator to provide unbiased estimates. As in the previous analysis in patients, the bifactor solution was compared with other solutions (i.e., unidimensional, multidimensional, and hierarchical models) using Log-Likelihood (LL), Akaike Information Criterion (AIC), Bayesian Information Criterion (BIC), and Sample-size Adjusted BIC (SABIC) as model fit statistics. Path diagrams that illustrate these models are presented in Supplementary Figure S1. Reliability and strength indices such as McDonald’s omega (ω_H_) (34), omega hierarchical (am) (34), and index H (35), were computed to determine: 1) the proportion of common variance accounted by general and specific symptom dimensions; 2) the proportion of reliable variance accounted by the general dimension not unduly affected by the specific dimensions; 3) the proportion of reliable variance accounted for by each specific dimension not unduly affected by the general and all the other specific dimensions; 4) the overall reliability and replicability of the bifactor construct of psychosis-like experiences. Finally, we generated factor scores for one general psychotic experience dimension and three specific dimensions of positive, negative, and depressive psychotic experiences.

For patients, we used the previously generated factor scores for one general psychosis dimension and five specific dimensions of positive, negative, disorganised, manic, and depressive symptoms (23).

### Symptom dimensions and cannabis use

We evaluated the relationship between psychotic symptom dimensions in patients, or psychotic experience dimensions in controls, and cannabis use using linear mixed effects models in STATA 14. We specifically modelled symptom dimension scores as a function of each of the six measures of cannabis use. We then evaluated the combined effect of frequency of use and potency of cannabis. To account for the non independence of symptom profiles of subjects assessed within the same country (for example, due to cultural similarities), and for the potential within-site correlation (for example, due to context factors), we fitted a three-level mixed model, where the random effect encompassed a double level of random intercepts: one due to the countries, and another due to the sites within the countries.

We conducted a sensitivity analysis to estimate an additional bifactor model based on the distress elicited by the psychotic experiences. This aimed to rule out the possibility in controls that 1) positive psychotic experiences were no distressing if associated to cannabis use, and 2) self-rating of positive psychotic experiences was influenced by contextual or cultural factors. We therefore confirmed our findings further adjusting the linear mixed effects models for the distress factor scores.

## Results

### Sample characteristics

We analysed data from 901 FEP patients and 1,235 controls. Main Sociodemographic characteristics and history of substance misuse of patients and controls are presented in Supplementary Table S1. Supplementary Tables S3.1 and S3.2 show the sample prevalence of psychotic experiences in controls and of psychotic symptoms in patients.

### Bifactor model of psychotic experiences in controls

Table 1 shows that, similar to our previous analysis of the OPCRIT items (23), the bifactor model provided the best fit for the CAPE items, as illustrated by AIC, BIC and SABIC substantially lower compared with competing models. This solution explained 60% of the unique variance. In addition, Figure 1 shows that, within the bifactor model, the explained variance was due to individual differences mostly on the general psychotic experience dimension. This is illustrated by the relative omega coefficient, which, for example, showed that 85% of the reliable variance was due to the general dimension when partitioning out the variability in scores due to the specific dimensions. Moreover, factor loadings of moderate to high magnitude were observed for most items on the general psychotic experience dimension, whereas factor loadings of a smaller magnitude were observed for the specific dimensions (Figure 1). Consistently, the index H, which is a measure of the construct reliability and replicability across studies (35), was veiy high for the general dimension (0.92), moderate for positive (0.78) and negative (0.71) dimensions and lower for the depressive dimension (0.41).

**Figure 1.**
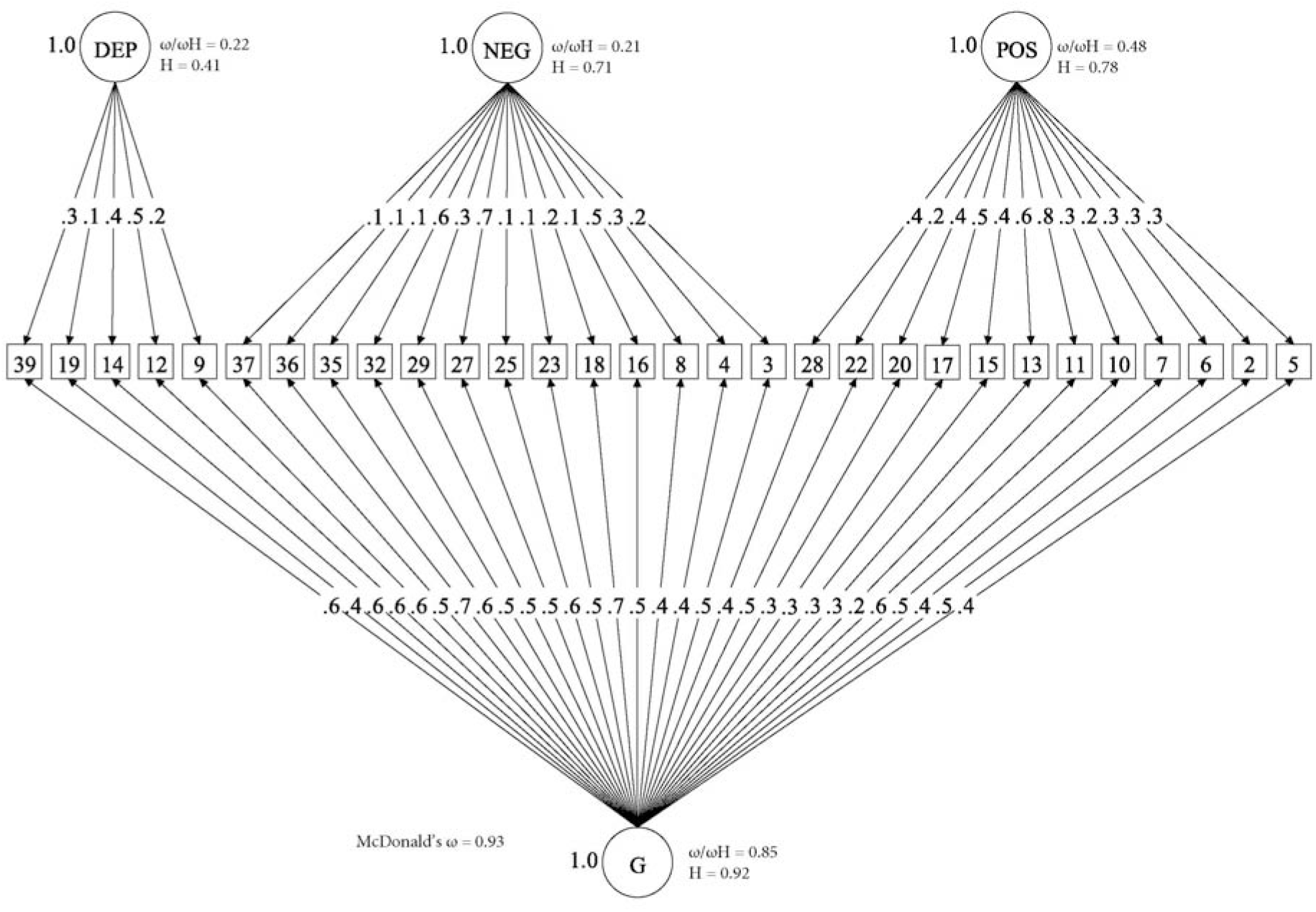
Bifactor model of psychotic experiences in controls. (□) Observed variables (No. of CAPE items); (○) Unobserved variables (latent iactors); (-») standardized item loading estimation onto latent iactors; G, general psychosis-like factor; Specific psychotic experiences iactors: DEP, Depression; NEG, Negative; POS, Positive. Reliability and strenght estimates: H=construct reliability index; ω= McDonald omega; ωH=hierarchical omega; *ω/ωπ=* Relative omega. Explanatory note: McDonald’s ω is an estimate of the proportion of the common variance accounted by general and specific symptom dimensions.(34) Relative omega (ω/ωh) is the amount of reliable variance explained in the observed scores attributable to a) the general factor independently from the specific symptom dimensions, and 2) each specific symptom dimension independently from the general factor. *H* is an index of the quality of the measurement model based on the set of CAPE items for each dimension.(35) Indices can range from 0 to 1, with values closer to 1 indicating a better construct reliability and replicability across studies.

**Table 1.** Model fit statistics of unidimensional, multidimensional, bifactor, second-order models for psychotic symptoms and psychotic experiences

| CAPE (CONTROLS) |  |  |  |  |
| --- | --- | --- | --- | --- |
|  | Full information fit statistics<sup>a</sup> |  |  |  |
|  | LL | AIC | BIC | SABIC |
| A - Unidimensional Model | -23638 | 47397 | 47715 | 47524 |
| B - Multidimensional Model (five uncorrelated factors) | -23844 | 47808 | 48126 | 47936 |
| C - Multidimensional Model (five correlated factors) | -23341 | 46808 | 47142 | 46942 |
| **D - Bifactor Model (one general factor and five specific uncorrelated factors)** | **-23139** | **46458** | **46935** | **46649** |
| E - Hierarchical Model (five first-order specific correlated factors and one second order general factor) | -23341 | 46807 | 47135 | 46938 |
| OPCRIT (PATIENTS) |  |  |  |  |
|  | Full information fit statistics<sup>a</sup> |  |  |  |
|  | LL | AIC | BIC | SABIC |
| A - Unidimensional Model | -29965 | 60126 | 60618 | 60306 |
| B - Multidimensional Model (five uncorrelated factors) | -28070 | 56335 | 56826 | 56515 |
| C - Multidimensional Model (five correlated factors) | -27894 | 56004 | 56546 | 56202 |
| **D - Bifactor Model (one general factor and five specific uncorrelated factors)** | **-27597** | **55489** | **56226** | **55759** |
| E - Hierarchical Model (five first-order specific correlated factors and one second order general factor) | -27995 | 56197 | 56713 | 56386 |

LL, log-likelihood; AIC, Akaike Information Criterion; BIC, Bayesian Information Criterion; SABIC
Sample-size Adjusted Bayesian Information Criterion
A difference of 10 in AIC, BIC and SABIC is considered important. Lower values indicate a statistically better model fit (best values across models are indicated in bold).

### Symptom dimensions in patients by lifetime cannabis use, current cannabis use, age at first use, and money spent on cannabis

Models’ results are presented in Table 2.1 which shows that:

1. There were no differences in the distribution of positive symptoms according to ever use of cannabis or early age at first use (=<15 years old). However, positive symptoms were more common among patients who currently used cannabis (B=0.22; 95%CI 0.04 to 0.4; p=0.016) and who spent more than 20 euros per week on cannabis (B=0.35; 95%CI 0.14 to 0.55; p=0.001).
2. Fewer negative symptoms were observed among those patients who used cannabis at least once compared with those who never tried (B = –0.27; 95%CI −0.42 to −0.12; <0.001). Early age at first use and current use of cannabis was not associated with negative symptomatology.
3. Manic symptoms were more frequent among patients who had either ever use (B=0·31; 95%CI 0.14 to 0.47; p<0.001) or current use of cannabis (B=0.19; 95%CI 0. 01 to 0.36; p=0.035)·
4. There were no differences in the distribution of the scores on the depressive and general psychosis dimensions according to any measure of cannabis use.

**Table 2.1.**
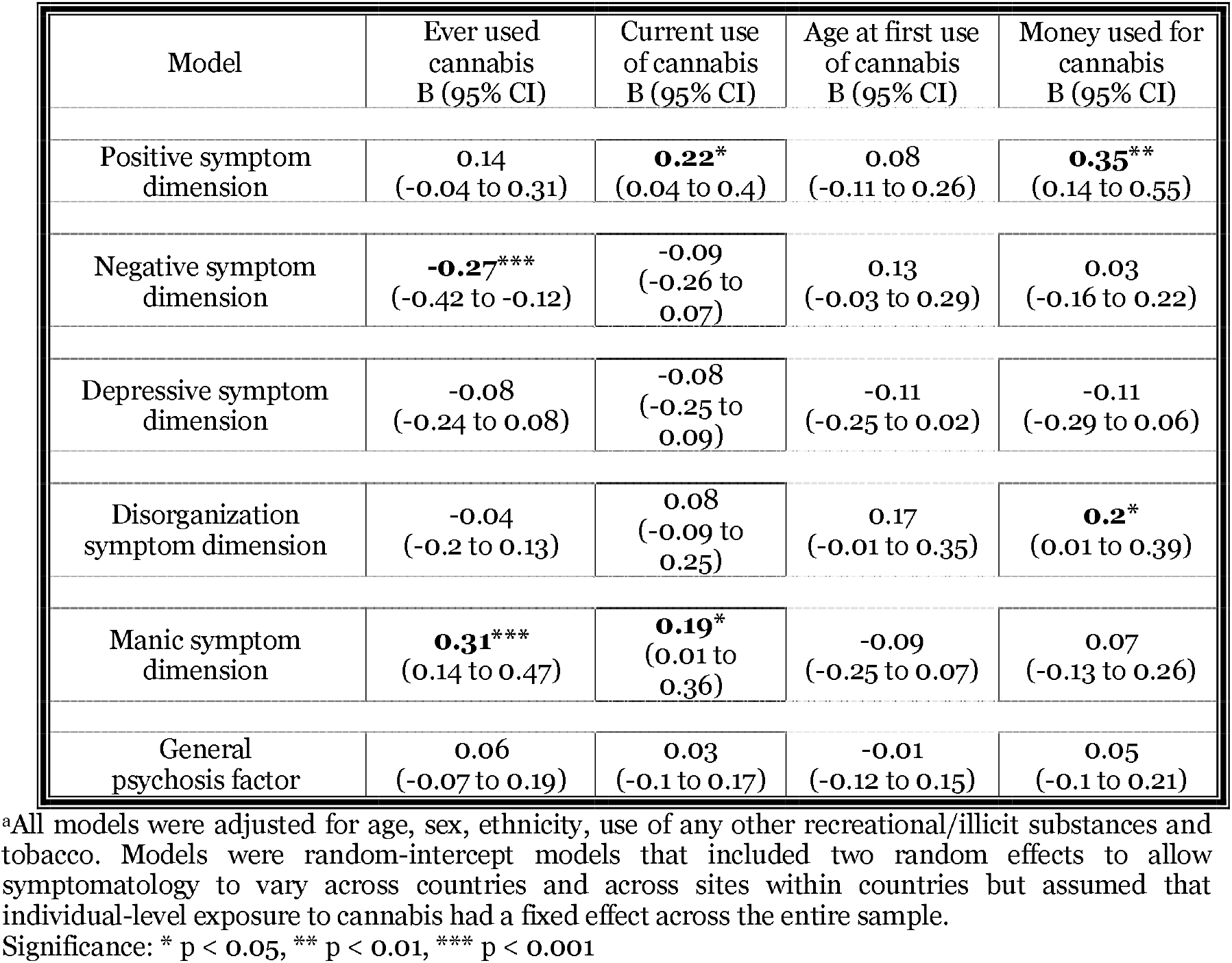
Symptom dimensions in FEP patients by measures of cannabis use^a^

| Model | Ever used cannabis<br>B (95% CI) | Current use of cannabis<br>B (95% CI) | Age at first use of cannabis<br>B (95% CI) | Money used for cannabis<br>B (95% CI) |
| --- | --- | --- | --- | --- |
| Positive symptom dimension | 0.14<br>(-0.04 to 0.31) | **0.22\***<br>(0.04 to 0.4) | 0.08<br>(-0.11 to 0.26) | **0.35\*\***<br>(0.14 to 0.55) |
| Negative symptom dimension | **-0.27\*\*\***<br>(-0.42 to -0.12) | -0.09<br>(-0.26 to 0.07) | 0.13<br>(-0.03 to 0.29) | 0.03<br>(-0.16 to 0.22) |
| Depressive symptom dimension | -0.08<br>(-0.24 to 0.08) | -0.08<br>(-0.25 to 0.09) | -0.11<br>(-0.25 to 0.02) | -0.11<br>(-0.29 to 0.06) |
| Disorganization symptom dimension | -0.04<br>(-0.2 to 0.13) | 0.08<br>(-0.09 to 0.25) | 0.17<br>(-0.01 to 0.35) | **0.2\***<br>(0.01 to 0.39) |
| Manic symptom dimension | **0.31\*\*\***<br>(0.14 to 0.47) | **0.19\***<br>(0.01 to 0.36) | -0.09<br>(-0.25 to 0.07) | 0.07<br>(-0.13 to 0.26) |
| General psychosis factor | 0.06<br>(-0.07 to 0.19) | 0.03<br>(-0.1 to 0.17) | -0.01<br>(-0.12 to 0.15) | 0.05<br>(-0.1 to 0.21) |

<sup>a</sup>All models were adjusted for age, sex, ethnicity, use of any other recreational/illicit substances and tobacco. Models were random-intercept models that included two random effects to allow symptomatology to vary across countries and across sites within countries but assumed that individual-level exposure to cannabis had a fixed effect across the entire sample.
Significance: \* $p < 0.05$ , \*\* $p < 0.01$ , \*\*\* $p < 0.001$

### Psychotic experiences’ dimensions in population controls by lifetime cannabis use, current cannabis use, age at first use, and money spent on cannabis

Models’ results are presented in Table 2.2, which shows that:

1. There were no differences in the distribution of positive psychotic experiences according to ever use of cannabis or early age at first use (=<15 years old). However, positive psychotic experiences were more commonly self-reported by subjects who currently used cannabis (B=0.19; 95%CI 0.01 to 0.37; p=0.037) and who spent more than 20 euros per week on cannabis (B=0.31; 95%CI 0.01 to 0.6; p=0.044).
2. There were no differences in the distribution of the depressive and negative experiences in population controls according to cannabis use.
3. The general psychotic experience dimension was associated with current use of cannabis (B=0.21; 95%CI 0.04 to 0.41; p=0.03).

**Table 2.2.**
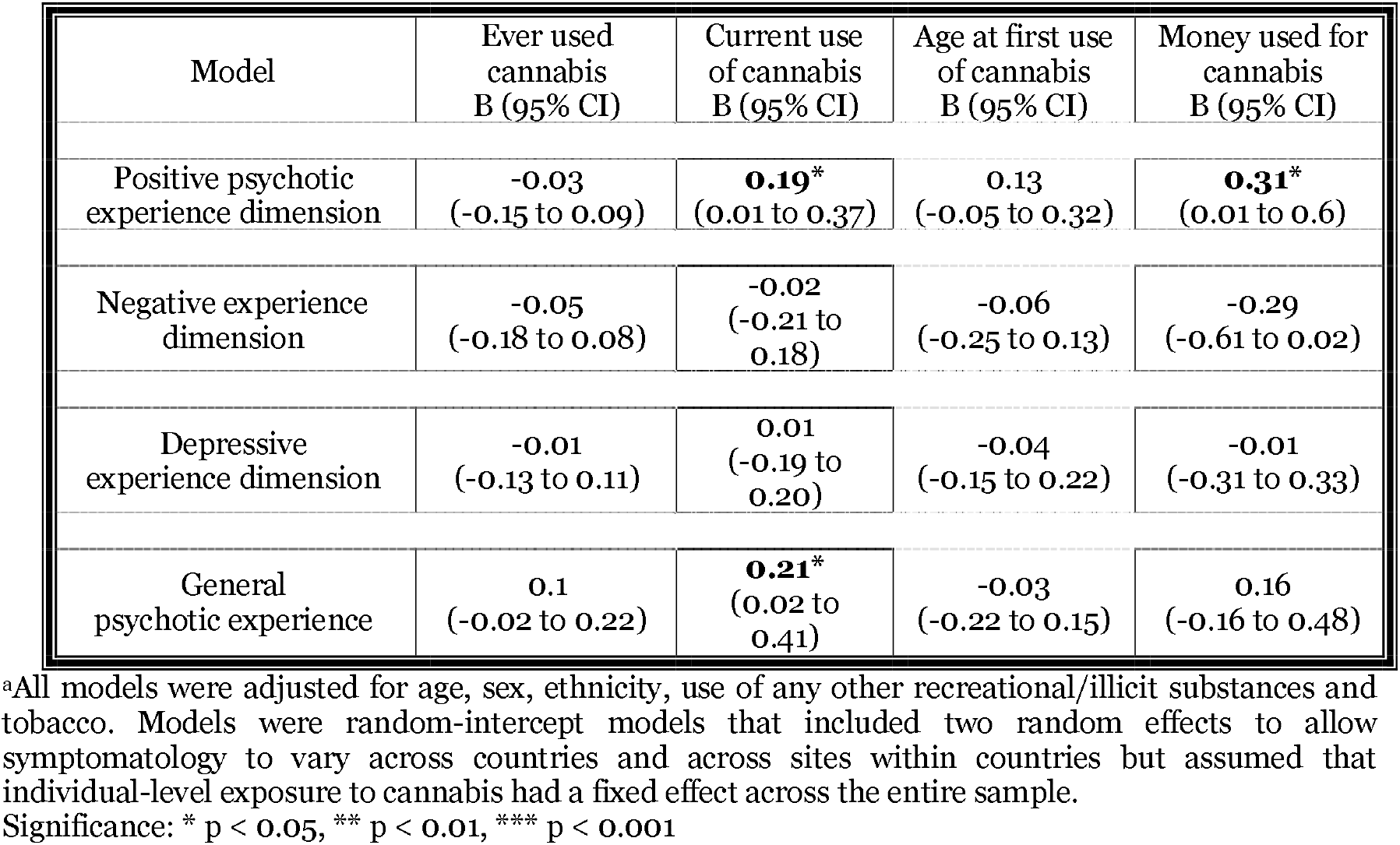
Psychotic experience dimensions in controls by cannabis use^a^

| Model | Ever used cannabis<br>B (95% CI) | Current use of cannabis<br>B (95% CI) | Age at first use of cannabis<br>B (95% CI) | Money used for cannabis<br>B (95% CI) |
| --- | --- | --- | --- | --- |
| Positive psychotic experience dimension | -0.03<br>(-0.15 to 0.09) | **0.19\***<br>(0.01 to 0.37) | 0.13<br>(-0.05 to 0.32) | **0.31\***<br>(0.01 to 0.6) |
| Negative experience dimension | -0.05<br>(-0.18 to 0.08) | -0.02<br>(-0.21 to 0.18) | -0.06<br>(-0.25 to 0.13) | -0.29<br>(-0.61 to 0.02) |
| Depressive experience dimension | -0.01<br>(-0.13 to 0.11) | 0.01<br>(-0.19 to 0.20) | -0.04<br>(-0.15 to 0.22) | -0.01<br>(-0.31 to 0.33) |
| General psychotic experience | 0.1<br>(-0.02 to 0.22) | **0.21\***<br>(0.02 to 0.41) | -0.03<br>(-0.22 to 0.15) | 0.16<br>(-0.16 to 0.48) |

<sup>a</sup>All models were adjusted for age, sex, ethnicity, use of any other recreational/illicit substances and tobacco. Models were random-intercept models that included two random effects to allow symptomatology to vary across countries and across sites within countries but assumed that individual-level exposure to cannabis had a fixed effect across the entire sample.
Significance: \* $p < 0.05$ , \*\* $p < 0.01$ , \*\*\* $p < 0.001$

### Symptom dimensions by lifetime pattern of cannabis use

Preliminary analysis testing independently the effects of frequency of use and potency of cannabis showed that, only in patients, positive symptoms were more common in those who used cannabis on a daily basis or exposed to high potency varieties (Supplementary Figure S2, Supplementary Table S4.1).

Testing the combined ‘type-frequency’ variable in patients, we found evidence of a linear relationship between the positive symptom dimension and the extent of exposure to cannabis, with daily users of high potency cannabis showing the highest score (B=0·35; 95%CI 0.14 to 0.56; p=0.001). Therefore, we introduced a contrast operator and plot the exposure-response relationship for positive symptoms (Figure 2), by comparing the predictive margins of the adjusted mean of each group against the grand adjusted mean of all groups. Figure 2 shows that the adjusted mean for daily users of high potency cannabis was 0.15 units greater than the grand adjusted mean. Moreover, the adjusted means for the groups who never or rarely used cannabis were respectively 0.13 or 0.14 units lower than the grand adjusted mean.

**Figure 2.**
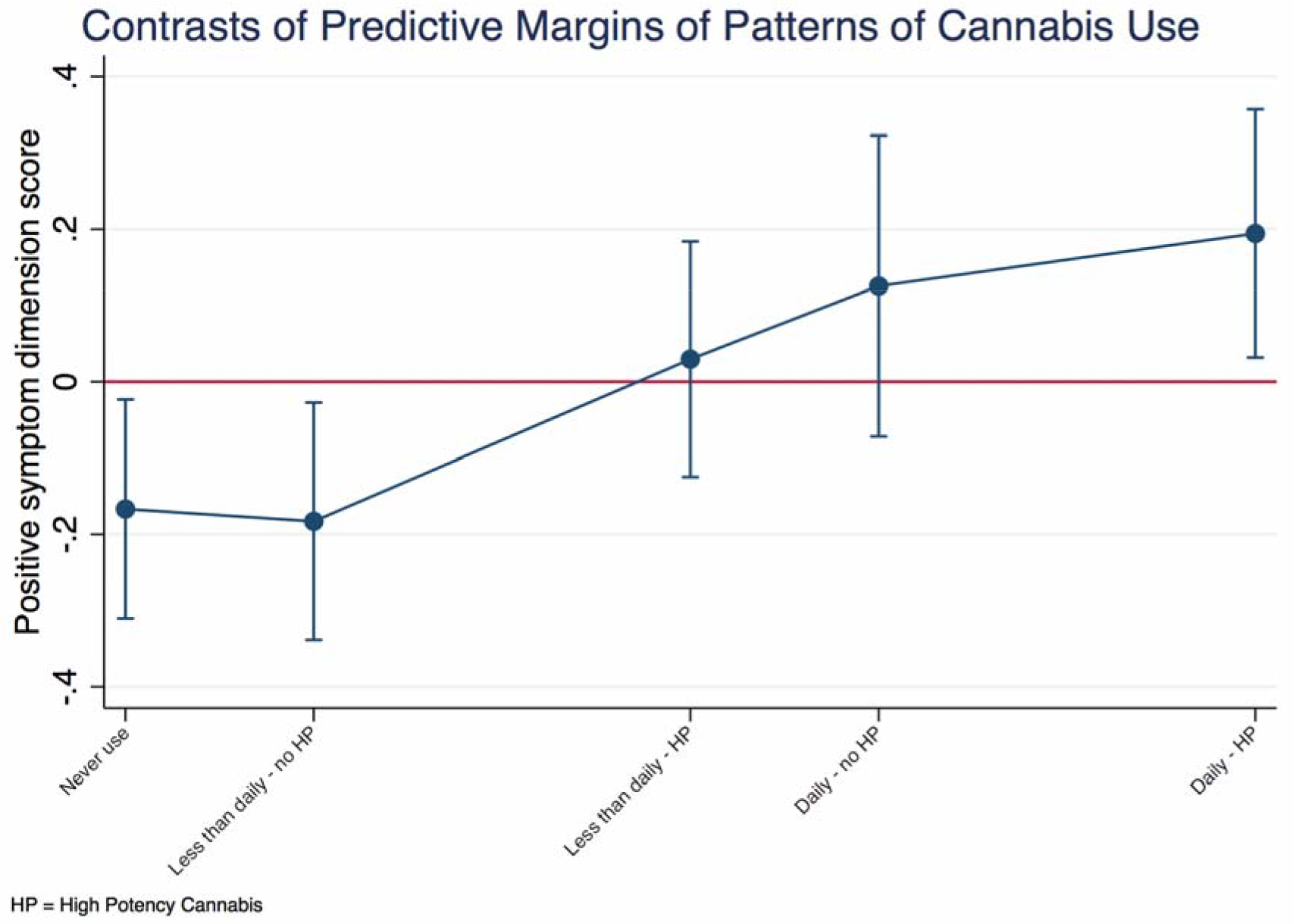
Positive symptom dimension in FEP patients by patterns of cannabis use. Figure 2 shows the contrasts of the positive symptom dimension predicted mean of each group against the predicted grand mean of all groups. Values were adjusted for age, sex, ethnicity, current use of other recreational and illicit substances (such as stimulants, hallucinogens, cocaine, crack, novel psychoactive substances) and tobacco. The model was a random intercept model which allowed symptoms to vary across countries and sites within countries but assumed frequency of use and type of cannabis had an individual fixed effect. The positive value for the contrast of the daily use of high potency cannabis indicates more positive symptomatology in this group. On the other hand, negative values for the contrasts of the first two groups indicates less positive symptomatology when there is less exposure to cannabis. These differences are statistically significant, as indicated by 95% confidence intervals that do not overlap with zero.

A negative relationship between the negative symptom dimension score and patterns of cannabis use was also observed in patients. Figure 3 shows that patients with psychosis who never used cannabis had more negative symptoms either compared with the grand adjusted mean or with any pattern of cannabis use.

**Figure 3.**
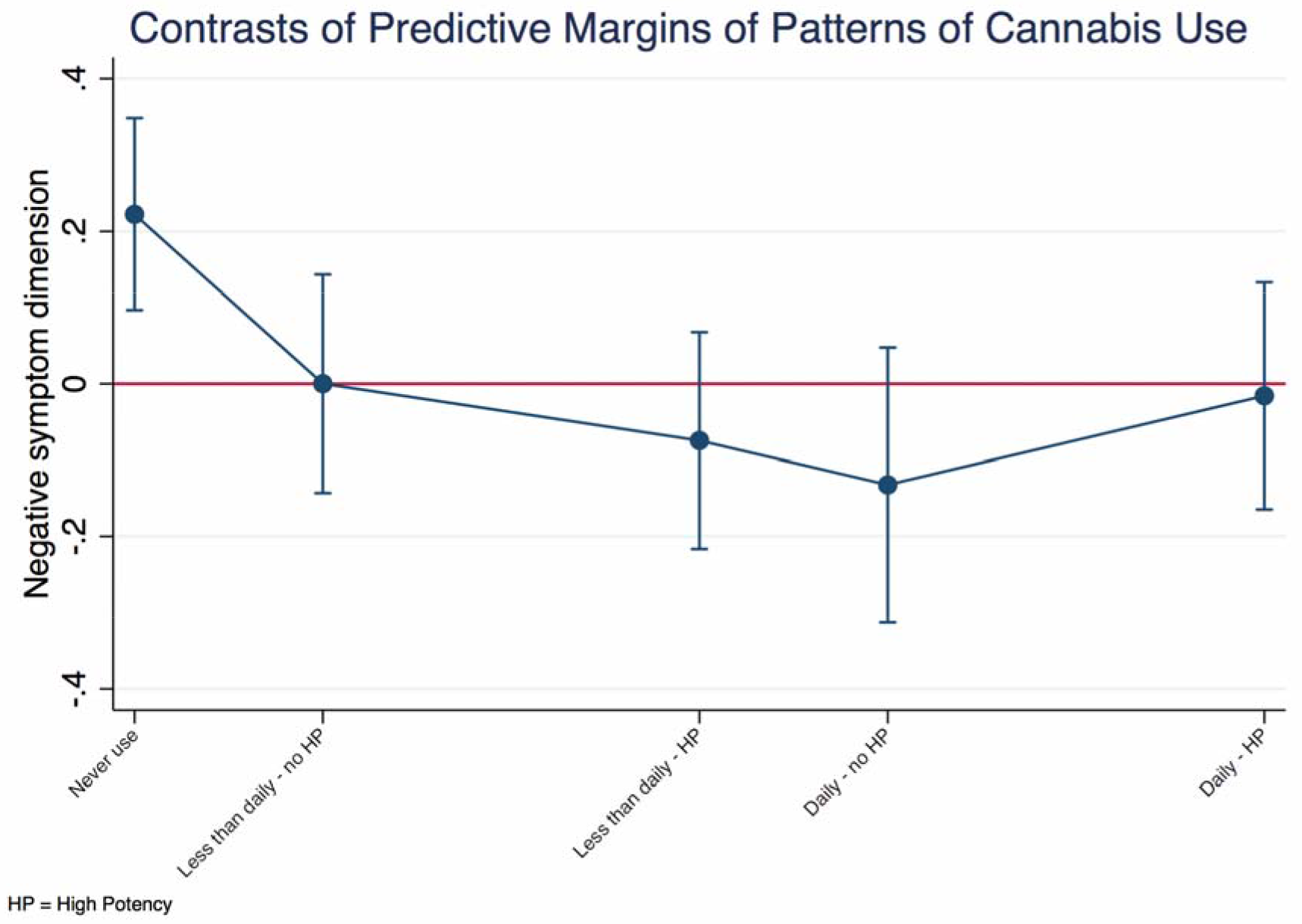
Negative symptom dimension by patterns of cannabis use. Figure 3 shows the contrasts of the negative symptom dimension predicted mean of each group against the grand adjusted predicted mean (represented by the red line). Subjects who had never used cannabis presented with more negative symptoms compared to the whole sample. Values are adjusted for age, sex, ethnicity, current use of other recreational and illicit substances and tobacco. The model was a random intercept model which allowed symptoms to vary across countries and sites within countries, but it assumed that frequency of use and type of cannabis had an individual fixed effect.

## Discussion

### Principal findings

This is the first multinational study exploring the dose effect of cannabis use on dimensions of symptom in FEP patients, and on dimensions of psychotic experiences in population controls. Our findings indicate that: 1) in patients, a positive correlation exists between the extent of premorbid cannabis use and the score at the positive symptom dimension, with daily users of high potency cannabis showing the worst symptomatology at FEP; 2) psychotic experiences in non-clinical populations is associated with current use of cannabis but is independent of the extent of lifetime exposure to cannabis; 3) negative symptoms at FEP are more common in patients who have never tried cannabis; 4) depressive symptoms are independent of any pattern of use of cannabis.

### Limitations

Our findings must be considered in the context of three main limitations. First, individual data on patterns of cannabis use are not validated with biological samples. However, such methods would not allow one to ascertain the extent of cannabis use over the years (36). Moreover, studies combining self-report and laboratory data support the reliability of subjects in reporting the type of cannabis they use (37, 38). Second, we did not take into account the cannabidiol (CBD) contribution to the potency variable, as official data on its content in the different cannabis varieties were not available in most study sites. Indeed, CBD might counterbalance Δ9-THC effects and minimise both psychotic experiences (39) and positive psychotic symptoms (40). Third, it could be argued that self-report checklists for subclinical symptoms, such as the CAPE, might yield false positive rates, as the interpretation and rating of the experiences might be subject to contextual or cultural factors in individuals without a psychiatric history (41). However, our findings are confirmed even after adjusting the analyses for the distress elicited by psychotic experiences, which is less subject to misinterpretation (42).

### Comparison with previous research

We extend previous research on cannabis and psychotic symptoms to a multinational sample confirming the association between cannabis use and positive symptoms of FEP (8, 9). Our results are in line with Schoeler et al. (2016), who carefully scrutinised the literature on the effect of continuation of cannabis use after FEP, concluding that this would be associated with a more severe positive symptomatology (43). That said, any comparison with previous research is limited by the lack of information on frequency and potency in all the previous studies along with subjects’ exposure to more potent varieties of cannabis over recent years (22).

In this respect, we firstly provide some evidence that cannabis affects positive symptoms in a dose response manner, further supporting the converging epidemiological and experimental evidence that the use of cannabis with high content of Δ9-THC has a more detrimental effect than other varieties (20, 29, 44).

We also report evidence in a multinational FEP sample of an association between lifetime cannabis use and fewer negative symptoms, the latter often considered as a marker of greater neurodevelopmental impairment in psychotic subjects. Indeed, our findings are consistent with the hypothesis that psychotic disorders could differ in their degree of neurodevelopmental impairment, thus being characterized, for example, by less neurodevelopmental features when associated with cannabis use (45). Support for this hypothesis is provided by evidence that additional markers suggestive of neurodevelopmental impairment in psychotic disorders tend to be uncommon in cannabis-associated psychosis (45).

On the other hand, some authors have suggested that people with a psychotic disorder might use cannabis as an attempt to self-medicate negative symptoms, thus the observed reduction in negative symptomatology would be an epiphenomenon due to the cannabis intake itself (16). In principle, this possibility is unlikely since a prominent negative symptom is reduction of social and instrumental skills, which are necessary to obtain cannabis and sustain its use over time. Further, our analysis indicates a lack of a dose-response relationship between cannabis use and the negative symptom dimension score, suggesting that the difference holds between those who never tried cannabis and those who may have used it only once. Finally, it could be argued that psychotic people use cannabis to self-medicate depressive symptoms, as opposed to negative symptoms. Again, this explanation is not supported by our study, which differentiates negative and depressive dimensions to disprove the relationship between depressive symptomatology and cannabis use.

Last, we report that the cumulative exposure to cannabis does not impact on psychotic experiences in controls. One could of course argue that the largest proportion of subjects with the harmful pattern of cannabis use were patients. However, further research is needed to look into plausible mechanisms of resilience to the psychotogenic effect of cannabis as observed in our controls, who report psychotic experiences if current users but do not seem to accumulate a risk over life time cannabis use and develop psychotic disorders. Indeed, future studies should aim to investigate if and how genetic factors, plausibly regulating the dopamine system, interact with cannabis use in shaping the quality of symptomatology.

### Implications

In this multinational study, we show that the pattern of use of cannabis explains part of the heterogeneity in positive and negative symptom dimensions at first episode psychosis. Further research should aim to determine biological mechanisms underlying how cannabis impacts on different clinical manifestations of psychosis. Meanwhile, translating current findings into clinical practice, symptom dimensions are useful to stratify patients and develop and implement secondary and tertiary prevention schemes for cannabis-associated psychosis. Treatment plans should integrate harm reduction and cannabis use cessation strategies, combined with possibly better tailored treatment strategies such as the adjunct to conventional pharmacological treatment of cannabidiol (40).

## Acknowledgment

The authors have no conflicts of interest to declare in relation to the work presented in this paper.

The EU-GEI Project is funded by the European Community’s Seventh Framework Programme under grant agreement No. HEALTH-F2-2010-241909 (Project EU-GEI). The Brazilian study was funded by the São Paulo Research Foundation under grant number 2012/0417-0. The work was further supported by: Clinician Scientist Medical Research Council fellowship (project reference MR/M008436/1) to MDF; Veni grant from the Netherlands Organisation for Scientific Research (grant no. 451-13-022) to UR. Funders were not involved in design and conduct of the study; collection, management, analysis and interpretation of the data; preparation, review or approval of the manuscript, and decision to submit the manuscript for publication.

This paper represents independent research part-funded by the National Institute for Health Research (NIHR) Biomedical Research Centre at South London and Maudsley NHS Foundation Trust and King’s College London. The views expressed are those of the author(s) and not necessarily those of the NHS, the NIHR or the Department of Health and Social Care.

